# A Genetically Encoded Far-Red Fluorescent Indicator for Imaging Synaptically-Released Zn^2+^

**DOI:** 10.1101/2022.06.02.494512

**Authors:** Tianchen Wu, Manoj Kumar, Shengyu Zhao, Mikhail Drobizhev, Xiaodong Tian, Thanos Tzounopoulos, Hui-wang Ai

## Abstract

Synaptic Zn^2+^ has emerged as a key neuromodulator in the brain. However, the lack of research tools for directly tracking synaptic Zn^2+^ in the brain in live animals hinders our rigorous understanding of the physiological and pathological roles of synaptic Zn^2+^. In this study, we developed a genetically encoded far-red fluorescent indicator for monitoring synaptic Zn^2+^ dynamics in the nervous system. Our engineered <u>f</u>ar-red fluorescent indicator for <u>s</u>ynaptic <u>Z</u>n^2+^ (FRISZ) displayed a substantial Zn^2+^-specific turn-on response and low micromolar affinity. We genetically anchored FRISZ to the mammalian extracellular membrane via a transmembrane α-helix. We further successfully used membrane-tethered FRISZ (FRISZ-TM) to image synaptic Zn^2+^ dynamics in response to sound in the primary auditory cortex (A1) in awake mice. This study thus establishes a new technology for studying the roles of synaptic Zn^2+^ in the nervous system.

## Introduction

The existence of labile zinc ions (Zn^2+^) and Zn^2+^-containing vesicles in the brain has been known for decades (*1-3*). Recent studies have clearly established the role of synaptically released Zn^2+^ in modulating neurotransmission (*4-12*). A significant subset of glutamatergic neurons can accumulate Zn^2+^ in synaptic vesicles via the vesicular Zn^2+^ transporter (ZnT3) and co-release Zn^2+^ with glutamate (*13, 14*). Synaptically released Zn^2+^ interacts with and modulates the activity of glutamate receptors, such as *N*-methyl-d-aspartate receptors (NMDAR) (*4, 9, 12*), α-amino-3-hydroxy-5-methyl-4-isoxazolepropionic acid receptor (AMPAR) (*5*), kainate receptors (KAR) (*15-17*); and GABA receptors (GABARs) (*11, 18-20*). In addition, synaptic Zn^2+^ may enter postsynaptic neurons via voltage-gated Ca^2+^ channels (VGCCs) or Ca^2+^-permeable AMPARs to trigger further signaling cascades (*3, 21*). Dysregulation of synaptic Zn^2+^ signaling has been linked to numerous neurological diseases, such as stroke, epilepsy, depression, Alzheimer’s disease, and hearing disorders (*22-24*). Although the presence and importance of labile Zn^2+^ in the nervous system have been widely appreciated, the dynamics of synaptic Zn^2+^ release in response to naturally occurring stimuli remain unclear. A further understanding of the physiological and pathological roles of synaptic Zn^2+^ in the nervous system requires convenient and robust tools to track synaptic Zn^2+^ dynamics directly in the brain in behaving animals and disease models.

Zn^2+^ is electrochemically inert and cannot be detected by electrochemical methods. Traditionally, biological Zn^2+^ is studied with histochemical procedures, such as dithizone-based colorimetric staining and Timm’s staining (*1, 25*). These invasive methods cannot be applied to living organisms and only provide an endpoint snapshot on Zn^2+^ localization. Over the past several decades, researchers have developed a number of fluorescent Zn^2+^ indicators, including small molecule sensors, genetically encoded sensors, and hybrid sensors (*26, 27*). Although synthetic and hybrid Zn^2+^ indicators have proven to be invaluable tools, it is challenging to use them for repeated measurements in live animals. Also, it is difficult to maintain a stable concentration of exogenously loaded fluorescent indicators for detecting secreted Zn^2+^ in the extracellular space in living organisms.

Alternatively, genetically encoded fluorescent indicators have shown promising results and gained much interest. They often display high specificity and are compatible with common signal peptides for convenient subcellular localization. Moreover, they can be readily delivered into live cells and organisms to achieve transient or long-term expression (*28, 29*). Furthermore, genetically encoded indicators are compatible with various viral vectors and genetically modified animals, allowing studies concerning specific cell types or brain regions. Finally, plasmids and viral vectors encoding genetically encoded indicators can be broadly disseminated and readily adopted by other interested researchers.

Existing genetically encoded Zn^2+^ indicators (GEZIs) are mainly based on the Zn^2+^-dependent modulation of Förster resonance energy transfer (FRET) between two fluorescent proteins (FPs) (*27, 28*). These ratiometric GEZIs have become powerful tools for imaging Zn^2+^ in intracellular space and organelles (*30, 31*). However, they have relatively large molecular sizes and broad fluorescence excitation and emission profiles, making it challenging to perform multicolor or/and multiplexed experiments. Furthermore, their modest dynamic range represents a hurdle for *in vivo* imaging applications. Thus, recent studies by us and others have developed single-FP-based, green fluorescent intensiometric GEZIs (*32-35*). Despite the progress, fluorescence imaging of synaptic Zn^2+^ secretion in the brain in live animals has not yet been achieved due to several remaining technical hurdles. First, the Zn^2+^ affinities of current GEZIs are typically in the picomolar range and too high for monitoring synaptic Zn^2+^ secreted from intracellular vesicles (*36*). Moreover, most GEZIs cannot be effectively routed to the exoplasmic membrane (*32*). Given that extracellularly released synaptic Zn^2+^ is organized into distinct spatial microdomains around the plasma membrane to modulate neurotransmission (*12*), the lack of extracellular-membrane-bound sensors has been a major obstacle to tracking synaptic Zn^2+^ dynamics *in vivo*.

The two exceptions are our previously developed ZnGreen2 and ZIBG2, which have nanomolar to low micromolar affinities suited for detecting Zn^2+^ secretion from intracellular vesicles (*32, 33*). Also, they use *Pyrococcus furiosus* Rad50 (*pf*Rad50) zinc hook peptides as the sensory elements, allowing a more effective localization of the GEZIs to the extracellular surface of mammalian cells via a platelet-derived growth factor receptor (PDGFR) transmembrane (TM) α-helix (*32, 33*). However, our previous attempts to use ZnGreen2 or ZIBG2 to image synaptic Zn^2+^ secretion in the brain in mice were unsuccessful. ZnGreen2 bestowed very poor photostability due to a reversible photoswitching of the chromophore, a phenomenon previously observed in photoswitchable FPs (*37*). Because ZnGreen2 is a Zn^2+^-dependent fluorescence-turn-off indicator, the photoswitching phenomenon made the interpretation of results unreliable. On the other hand, ZIBG2 has much-enhanced photostability, but its small dynamic range precludes its usefulness *in vivo (32*). Current studies on Zn^2+^ in the brain often rely on Ca^2+^ indicators (*e*.*g*., GCaMPs) (*38*) to monitor neuronal activities in response to perturbations with Zn^2+^-specific chelators (*e*.*g*., ZX1) (*4*) or elimination of synaptic zinc in ZnT3 knockout mouse (*14*), which are inconclusive for distinguishing direct versus circuit-dependent synaptic Zn^2+^ effects.

It is a long-sought-after goal to engineer genetically encoded indicators with excitation and emission in the 600-900 nm near-infrared (NIR) optical window. The red-shifted indicators would require lower-energy and less-damaging photons for excitation than common genetically encoded indicators based on green or red FPs (GFPs or RFPs). Moreover, they are expected to enhance imaging depth because of reduced light scattering, absorption, and autofluorescence of the animal tissue (*39*). Previously, near-infrared (NIR) Ca^2+^ indicators were developed from a biliverdin-binding NIR FP (*40, 41*), but the requirement of the biliverdin cofactor is an obvious limitation because biliverdin is involved in normal physiology (*42*) and possibly limited under many conditions (*43*). We recently converted a far-red FP (FrFP), mMaroon1 (*44*), which does not require biliverdin and can spontaneously form a chromophore within its peptide sequence, into several bright, circularly permuted (cpmMaroon) variants with excitation and emission maxima longer than 600 nm (*45*). Here, we describe the engineering of a novel far-red fluorescent indicator for <u>s</u>ynaptic <u>Z</u>n^2+^ (FRISZ) from a cpmMaroon variant and the application of FRISZ in imaging synaptic Zn^2+^ release in awake mice in response to sound stimuli.

## Results

### Engineering of a far-red fluorescent indicator for synaptic Zn^2+^ (FRISZ)

The Rad50 zinc hook can undergo Zn^2+^-mediated homodimerization (*46*). Built upon our previous progress in engineering green fluorescent GEZIs, we examined the fusion of the *pf*Rad50 zinc hook to our new cpmMaroon variants. To date, single-FP-based indicators are almost always based on circularly permuted FPs (cpFPs) with termini in β-strand 7, and the modulation of chromophore ionization by the fused sensory domains is considered crucial for the responsiveness of single-FP-based indicators (*29*). However, we were unable to generate fluorescent cpmMaroon variants with circular permutation sites in β-strand 7 of mMaroon1 (*45*). Fortunately, we still identified two cpmMaroon variants, namely cpmMaroon185-186 and cpmMaroon196-186, with circular permutation sites in the loop between β-strands 9 and 10 but still sensitive to pH changes around pH 7.4 (**Fig. S1**) (*45*). We genetically fused a copy of the zinc hook to each of the termini of cpmMaroon185-186 or cpmMaroon196-186. To our delight, grafting the zinc hooks to cpmMaroon185-186 resulted in a fusion protein showing a 40% fluorescence increase in response to Zn^2+^ (**Fig. 1 & Fig. S2**). Next, we randomized the two linkers between cpmMaroon185-186 and the zinc hook elements, and screening the libraries led to a two-fold improvement in Zn^2+^-dependent fluorescence responsiveness. Moreover, we carried out random mutagenesis and performed 11 rounds of directed evolution (**Figs. S2 & S3**), further enhancing the Zn^2+^ responsiveness by about eight-fold. Finally, we replaced the Cys residues in the zinc hook elements with His, because we aim to express the indicator at the cell surface and His can bind Zn^2+^ but is more resistant to oxidation than Cys.

**Fig. 1.**
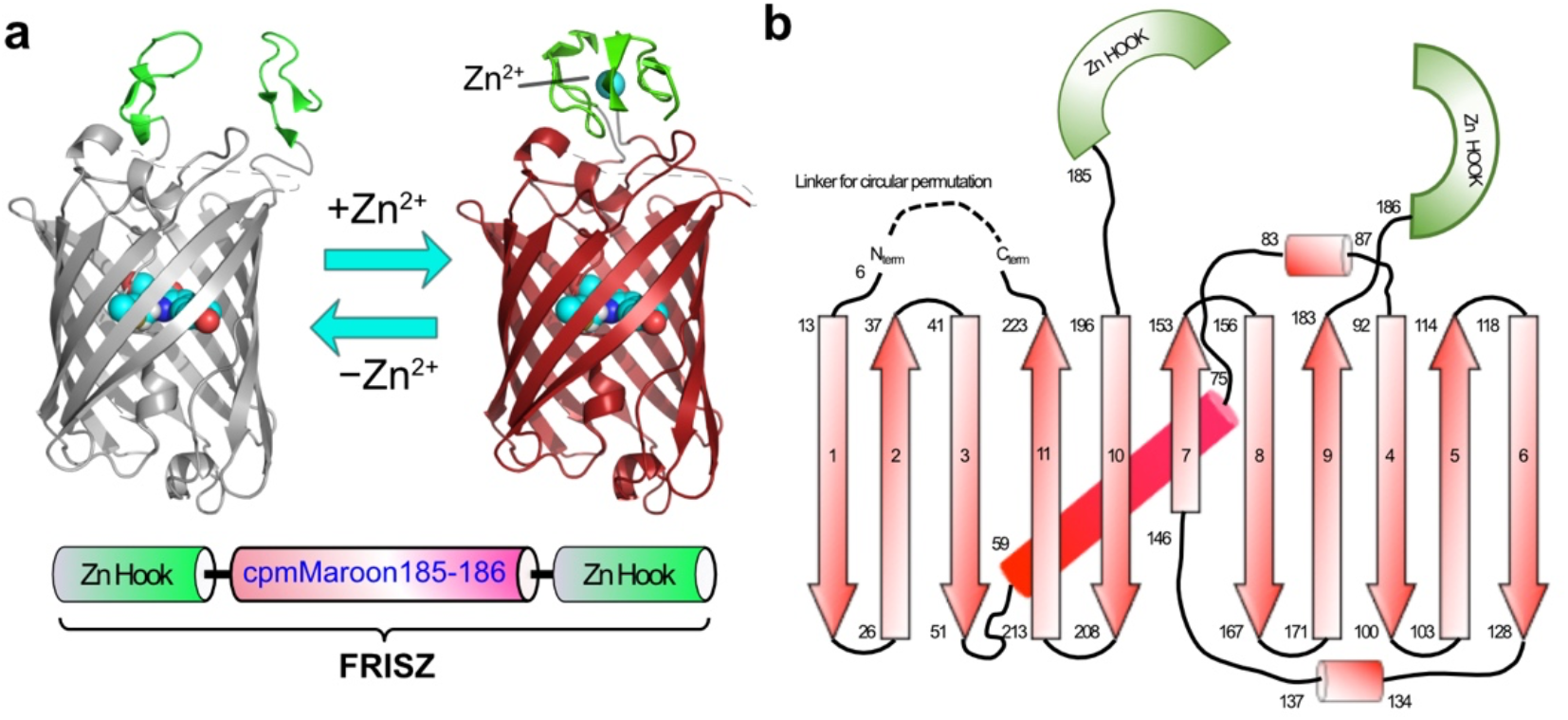
Illustration of the Zn^2+^-sensing mechanism and the secondary structure of FRISZ. **(a)** Illustration of FRISZ, in which two Rad50 zinc hook motifs are fused to the N- and C-termini of cpMaroon185-186. The domain arrangement of the primary sequence is also presented at the bottom. **(b)** Schematic representation of the secondary structure elements of FRISZ. Secondary structure elements in cpmMaroon185-186 are based on a mMaroon1 structure predicted by SWISS-MODEL using mCardinal (PDB: 4OQW) as the template. Cylinders represent α-helices, and arrows represent β-strands. The Rad50-derived zinc hook motifs are presented as green semicircles.

### Spectroscopic characterization

The final variant, termed FRISZ, exhibited a Zn^2+^-dependent 7.5-fold (F/F_0_) fluorescence-turn-on response (**Fig. 2a**). Its one-photon (1P) fluorescence excitation and emission maxima were ∼ 607 and 650 nm, respectively. When adding Zn^2+^ to the protein, we observed an increase in the absorption at ∼ 607 nm and a simultaneous absorption decrease at ∼ 460 nm (**Fig. 2b**), suggesting a Zn^2+^-dependent chromophore deprotonation process. Moreover, the Zn^2+^ addition enhanced the fluorescence quantum yield of the protein by ∼ 2-fold (**Table 1**). Thus, both the chromophore ionization and fluorescence quantum yield changes are responsible for the substantial turn-on response of FRISZ. In the Zn^2+^-bound state, the molecular brightness of FRISZ was ∼50% of mMaroon1 (**Table 1**). We further measured the fluorescence of FRISZ in the presence of various concentrations of free Zn^2+^ and determined the apparent dissociation constant (*K*_d_) to be ∼ 8.8 μM **(Fig. 1c**). In addition, we examined the specificity of FRISZ. Except for Zn^2+^, none of the tested cellularly abundant metal ions induced any notable fluorescence change (**Fig. 1d**). Under two-photon (2P) excitation conditions, FRISZ displayed excitation peaks at 1128 and 1216 nm (**Fig. 2e)**. Superior 2P brightness and Zn^2+^-induced fluorescence responses were obtained between 1050 and 1250 nm (**Table 1 & Fig. 2f**).

**Table 1.** Photophysical properties of FRISZ and other related proteins.

| Protein name | FRISZ |  | mMaroon1<sup>a</sup> | cpmMaroon185-186<sup>b</sup> |
| --- | --- | --- | --- | --- |
|  | Zn<sup>2+</sup>-free | Zn<sup>2+</sup>-bound |  |  |
| Peak Excitation $\lambda_{\text{ex}}$ (nm) | 607 | 606 | 609 | 610 |
| Peak Emission $\lambda_{\text{em}}$ (nm) | 650 | 654 | 657 | 648 |
| $F/F_0$ | 7.5 | | N/A<sup>c</sup> | N/A<sup>c</sup> |
| $K_d$ ($\mu\text{M}$) | 8.8 | | N/A<sup>c</sup> | N/A<sup>c</sup> |
| Apparent Hill coefficient ($n_H$) | 0.94 | | N/A<sup>c</sup> | N/A<sup>c</sup> |
| Fluorescence lifetime $\tau$ (ns) | 0.51 | 0.83 | N/D<sup>d</sup> | N/D<sup>d</sup> |
| Relative fraction of “anionic” chromophore $\rho_A$ | 0.19 | 0.49 | N/D<sup>d</sup> | N/D<sup>d</sup> |
| Effective extinction coefficient $\epsilon$ ($\text{mM}^{-1}\text{cm}^{-1}$)<sup>e</sup> | 11.5 | 32 | 80 | 54 |
| Quantum yield $\phi$ | 0.075 | 0.14 | 0.11 | 0.11 |
| 1P brightness ($\epsilon \times \phi$) | 0.86 | 4.5 | 8.8 | 5.9 |
| 2P brightness (GM) at the indicated excitation wavelength | 0.41 (1128 nm)<br>0.38 (1216 nm) | 2.4 (1124 nm)<br>2.2 (1208 nm) | N/D<sup>d</sup> | N/D<sup>d</sup> |
| $F/F_0$ at indicated 2P excitation wavelength | 6.5 (1064 nm)<br>6.1 (1124 nm)<br>5.9 (1208 nm) | | N/A<sup>c</sup> | N/A<sup>d</sup> |

<sup>a</sup> values adapted from Reference (44).
<sup>b</sup> values adapted from Reference (45).
<sup>c</sup> not applicable.
<sup>d</sup> not determined.
<sup>e</sup> presented as per mM of the total protein (a product of $\rho_A$ and the true extinction coefficient per mM of the anionic chromophore).

**Fig. 2.**
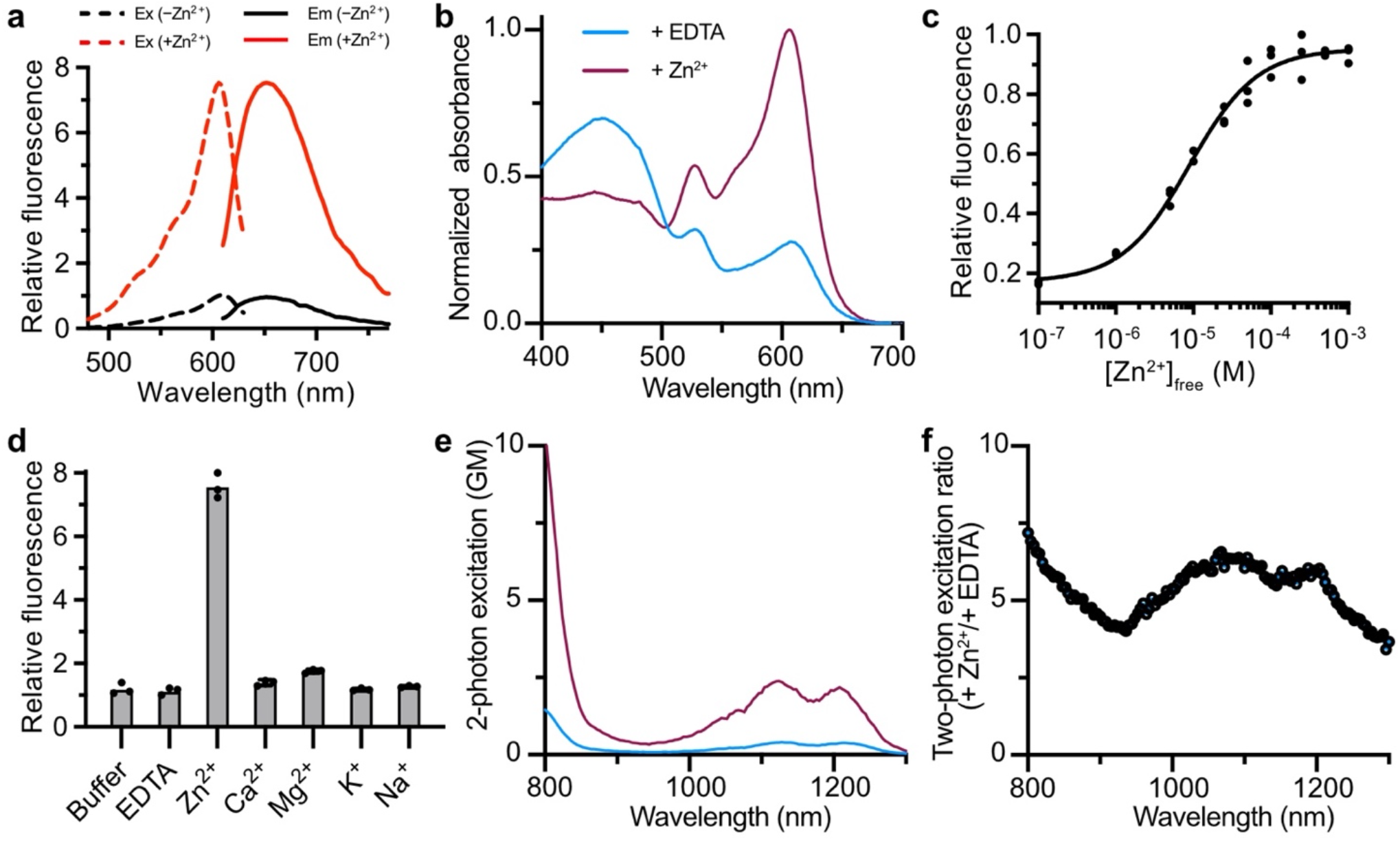
Spectroscopic characterization of FRISZ as a purified protein. **(a)** Excitation (dotted line) and emission (solid line) spectra of purified FRISZ in the presence of 100 μM Zn^2+^ (red) or 100 μM EDTA (black). **(b)** One-photon absorbance spectra of FRISZ in the presence of 100 μM Zn^2+^ or 100 μM EDTA. **(c)** Fluorescence of FRISZ in response to Zn^2+^ and other indicated chemicals (100 μM). n = 3. **(d)** Dose-response curve of purified FRISZ (*K*_d_ = 8.8 ± 1.0 μM). n = 3. **(e)** Two-photon fluorescence excitation spectra of FRISZ in the presence of 100 μM Zn^2+^ or 100 μM EDTA. **(f)** The ratio of two-photon excitation plotted against wavelength.

### Characterization of FRISZ at the mammalian cell surface

We next inserted the FRISZ gene into the commercial pDisplay plasmid, which encodes an N-terminal cleavable immunoglobin κ (Igκ) leader sequence for mammalian secretory pathway targeting and a C-terminal PDGFR TM domain for cell-surface anchoring (**Fig. 3a**). We observed excellent cytoplasmic membrane localization of FRISZ in transiently transfected human embryonic kidney (HEK) 293T cells (**Fig. 3a & Fig. S4**). Furthermore, Zn^2+^ addition triggered a robust fluorescence increase (F/F_0_ = ∼ 3) of cell-surface-localized FRISZ (FRISZ-TM) (**Fig. 3ab**). In contrast, we consistently observed no fluorescence response from cell-surface-localized mMaroon1 (mMaroon1-TM). The fluorescence increase of FRISZ-TM was dependent on Zn^2+^ concentrations, and the *K*_d_ of was determined to be ∼ 7.3 μM (**Fig. 3bc**). Moreover, we imaged the HEK 293T cells while locally applying Zn^2+^ via a micropipette. We determined the half-time (*t*_0.5_) for fluorescence rise (Zn^2+^ association) and decay (spontaneous dissociation) to be 117 ms and 1.28 s, respectively (**Fig. 3d**). Thus, the results collectively suggest that FRISZ-TM is promising for monitoring the dynamics of secreted Zn^2+^ from intracellular granules.

**Fig. 3.**
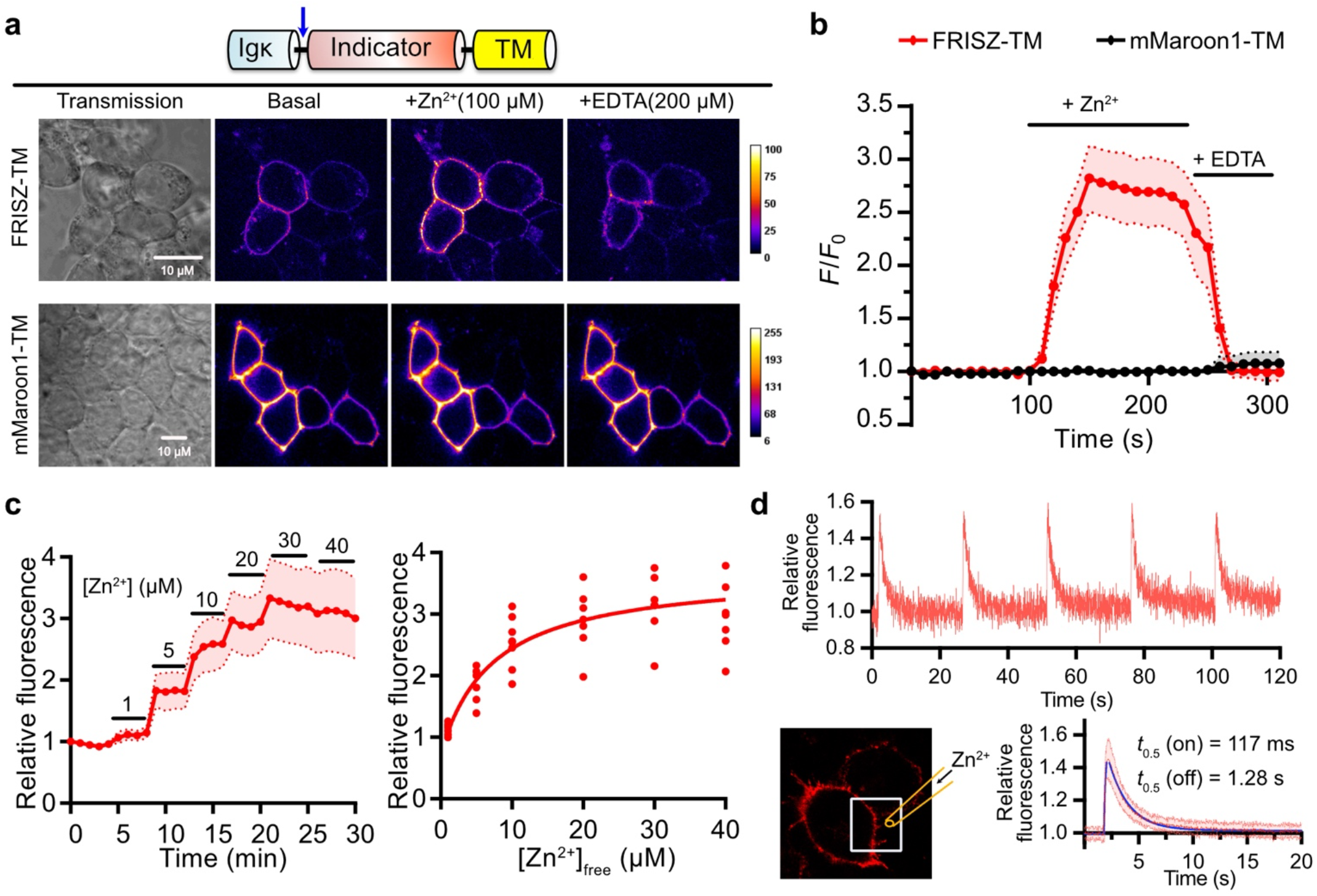
Characterization of FRISZ at the mammalian cell surface. **(a)** Top: Schematics of the genetic elements for the localization of FRISZ or mMaroon1 at the mammalian cell surface. Igκ, murine Igκ chain signal peptide. TM, PDGFRβ transmembrane domain. The arrow indicates the signal peptide cleavage site. Bottom: Representative pseudocolor images of FRISZ or mMaroon1 at the surface of HEK 293T in response to Zn^2+^ and EDTA. This experiment was repeated three times with similar results using independent cultures. Scale bar, 10 μm. **(b)** Quantification of time-lapse responses of FRISZ or mMaroon1 at the surface of HEK 293T cells to 100 μM Zn^2+^ and 200 μM EDTA. n=10 cells from three cultures. **(c)** Dose-response of FRISZ at the surface of HEK 293T (*K*_d_ = 7.3 ± 3.2 μM). Data are presented as mean and s.d. of 8 cells. **(d)** Response of FRISZ at the surface of HEK 293T to puff application of Zn^2+^. Top: Five sequential applications to a single cell. Bottom: Data presented as mean and s.d. of 30 repeats (six cells with five applications each) and fitted for monoexponential growth and decay.

### Use of membrane-tethered FRISZ to image synaptic Zn^2+^ in awake mice during audition

By using a fluorescent Ca^2+^ imaging approach in awake mice, our previous studies established that cortical synaptic Zn^2+^ signaling in the primary auditory cortex (A1) is elicited by natural sound stimuli and fine-tunes the sound frequency and response gain of the A1 neurons (*47, 48*). Moreover, in the absence of cortical synaptic Zn^2+^ (chelation with the high-affinity specific extracellular Zn^2+^ chelator, ZX1, or in ZnT3 knockout mice lacking synaptic Zn^2+^), mice exhibited reduced acuity for detecting changes in sound frequency (*48*), suggesting that synaptic Zn^2+^ contributes to sound-frequency tuning and enhances acuity for sound frequency discrimination. However, due to a lack of suitable Zn^2+^ indicators, we were previously unable to track synaptic Zn^2+^ dynamics in the A1 in live mice in response to sound stimuli. In this context, we examined the use of FRISZ-TM to detect synaptic Zn^2+^ release *in vivo* in response to sound stimuli. We prepared an adeno-associated viral (AAV) vector for the neuronal expression of FRISZ-TM (**Fig. 4a**). We next administered the FRISZ-TM AAV into the A1 of C57BL/6J mice and performed *in vivo* wide-field imaging 3-4 weeks post-viral injection (**Fig. 4ab**). We detected bright far-red fluorescence from the A1 of mice expressing either FRISZ-TM or the mMaroon1-TM control (**Fig. 4c**). Next, we presented broadband sounds to the awake mice and observed robust and reproducible sound-evoked fluorescence increases in FRISZ-TM-expressing mice (**Fig. 4de**). Infusion of ZX1 (*4, 49*) to the A1 effectively suppressed sound-evoked fluorescence changes, further confirming that the signal of FRISZ-TM is dependent on extracellular Zn^2+^. Furthermore, the mMaroon1-TM control mice showed negligible response to sound stimulation no matter whether ZX1 was present. These results support that FRISZ-TM allows for successful imaging of extracellular zinc increases *in vivo* in response to sound.

**Fig. 4.**
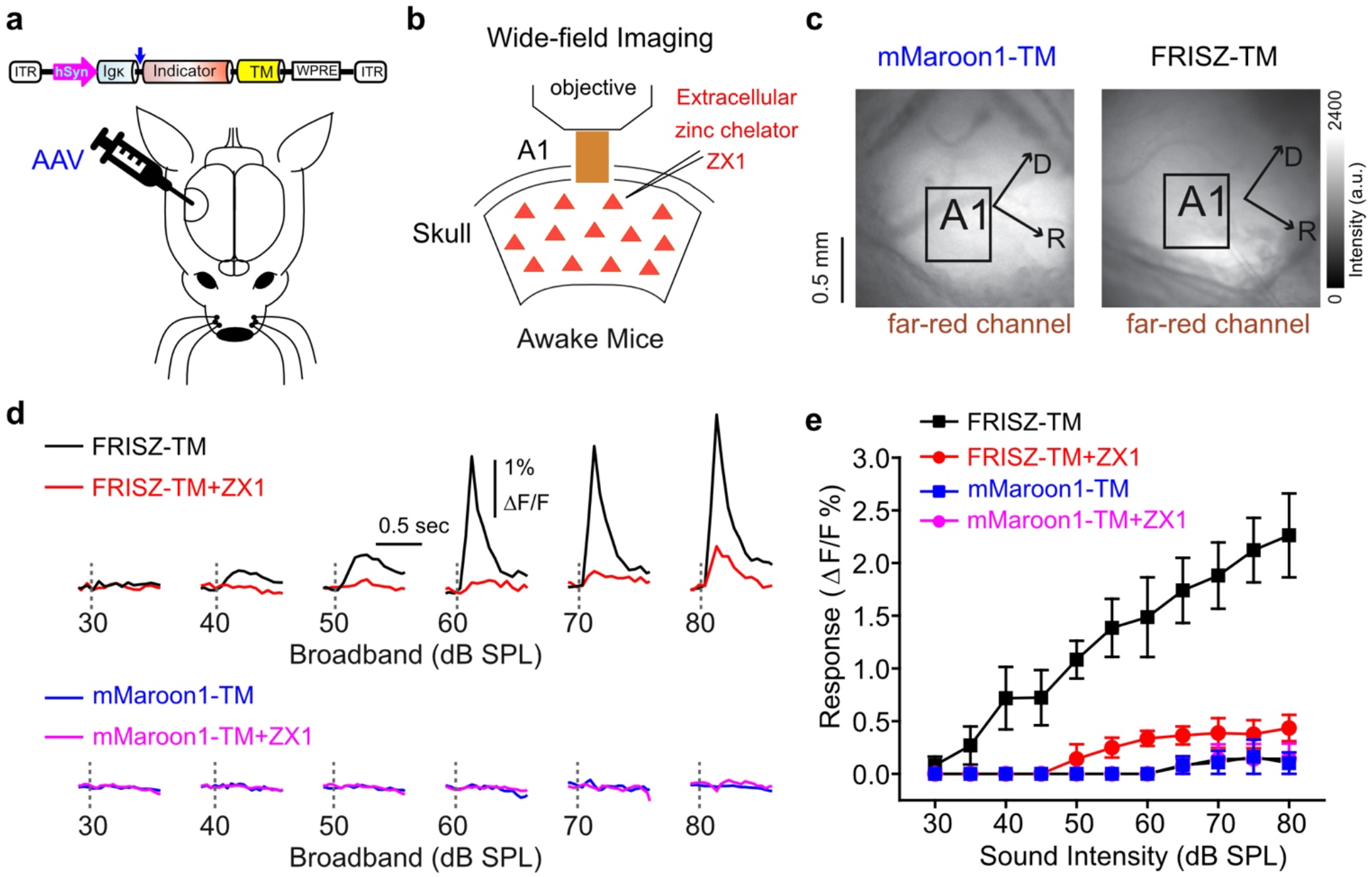
Imaging of synaptically released Zn^2+^ with cell-surface-bound FRISZ (FRISZ-TM) in the primary auditory cortex in head-fixed awake mice. **(a)** Schematics of the genetic elements of AAVs and viral injection into the auditory cortex in mice. **(b)** Illustration of transcranial imaging of cell-surface-localized FRISZ or mMaroon1 in head-fixed awake mice. Sounds are delivered through a calibrated speaker. The Zn^2+^-chelator, ZX1, was infused into the auditory cortex via a glass pipette connected to a syringe pump. **(c)** Fluorescence images of the primary auditory cortex (A1) showing the expression of mMaroon1-TM (left) and FRISZ-TM (right). D, Dorsal; R, rostral. **(d)** Top: Representative transcranial FRISZ fluorescence signals to broadband sounds before (black) and after ZX1 infusion (red). Bottom: Representative transcranial mMaroon1 fluorescence signals to broadband sounds before (blue) and after ZX1 infusion (magenta). **(e)** Summary plot of average sound-evoked fluorescence signals of FRISZ-TM (n = 5) and mMaroon1-TM (n = 3) mice. Error bars indicate s.e.m.

## Discussion

We have engineered the first far-red fluorescent GEZI by genetically fusing Rad50 zinc hook elements to cpmMaroon185-186, followed by multi-steps of directed evolution. FRISZ responded to low micromolar Zn^2+^ with excellent specificity and response magnitude. We further anchored FRISZ to the cell surface and successfully detected Zn^2+^ release in the A1 of awake mice in response to sound stimuli. Our results collectively suggest that FRISZ is a promising tool for studying Zn^2+^ release and Zn^2+^ plasticity during normal and pathological sensory processing.

We developed FRISZ from cpmMaroon185-186, which has new termini between β-strands 9 and 10 within the typical FP fold (**Fig. 1**). This new indicator has a different topology from other single-FP-based indicators, which typically use cpFPs with termini in β-strand 7 (*29*). One previous study reported several voltage indicators derived from mKate circularly permuted between β-strands 9 and 10, but the responses of those voltage indicators were mediocre (*50*). Thus, FRISZ provides a successful example of building highly responsive FP-based indicators in this new topology, and thus, we expect our result to inspire future sensor development.

FRISZ belongs to a new class of genetically encoded far-red fluorescence indicators, which are expected to reduce phototoxicity and autofluorescence and facilitate multiplexing and deeper tissue imaging. FRISZ exhibited peak excitation and emission above 600 nm and is among the most red-shifted FP-based indicators. A recent preprint described far-red fluorescent Ca^2+^ indicators based on circularly permuted FP, mKelly2 (*51*). Still, the excitation and emission of FRISZ are more red-shifted than these Ca^2+^ indicators by ∼ 10 nm, allowing the effective excitation of FRISZ with common ∼ 630 nm lasers or LEDs. We thereby expect that the cpmMaroon scaffold of FRISZ may be adapted to expand the color palette of genetically encodable indicators for other analytical targets.

FRISZ displayed strong far-red fluorescence under both 1P and 2P excitation. For 2P imaging, standard tunable Ti-sapphire lasers can excite FRISZ near 1060 nm to gain good brightness and Zn^2+^-responsiveness. Moreover, because the fluorescence of FRISZ under 1200 nm excitation is relatively strong (*52*) and there is an emerging interest in using 1200-nm excitation to increase brain tissue transmittance (*53*), FRISZ may be a suitable indicator for these future experiments.

FRISZ is the first GEZI successfully used for time-lapse imaging of Zn^2+^ secretion in the brain in live mice. However, since its responsiveness, particularly *in vivo*, is not yet optimal, we plan to perform further studies to enhance the robustness of FRISZ. Moreover, in addition to the brain, FRISZ may be used to monitor the secretion of vesicular Zn^2+^ in other tissue types, such as pancreatic islets (*32*).

Neuronal “tuning” to specific features of sensory stimuli is a fundamental property of sensory processing (*54-57*). In the A1, cortical neurons respond to selective ranges of sound frequencies. This tuning reflects the interplay of thalamocortical and intracortical inputs (*58*); however, the modulatory mechanisms that refine A1 tuning remain unclear. Synaptic Zn^2+^ is a potent neuromodulator that contributes to the fine-tuning of sound-frequency selectivity of A1 neurons (*11, 48*). Since here we have demonstrated the feasibility of using FRISZ-TM to monitor sound-evoked Zn^2+^ dynamics in the A1 in live mice during audition, our future studies will use the new tool to study cellular and circuit specificity of cortical synaptic Zn^2+^ dynamics and their effects on A1 neuronal sound processing. Furthermore, because synaptic Zn^2+^ is a potent neuromodulator throughout the cortex, our new sensor will improve the understanding of the roles of synaptic Zn^2+^ in cortical information processing beyond the A1 and beyond even sensory cortices (*3, 16, 59*).

## Materials and Methods

### Key materials and general methods

All chemicals were purchased from Fisher Scientific, Sigma-Aldrich, Thomas Scientific, and VWR. DNA oligos were purchased from IDT and Eurofins Genomics. Restriction enzymes were purchased from Thermo Scientific. DNA sequences were analyzed by Eurofins Genomics. Phusion High-Fidelity DNA polymerase from Thermo Fisher was used for DNA amplification and cloning. Taq DNA polymerase from New England Biolabs (in the presence of Mn^2+^) and GeneMorph II Random Mutagenesis Kit from Agilent were used for error-prone PCR. pcDNA3.1-mMaroon1 (*44*) (Addgene plasmid # 83840) was a gift from Michael Lin (Stanford). pCMV(MinDis). pCMV(MinDis)-iGluSnFR (*60*) (Addgene plasmid # 41732) was a gift from Loren Looger (HHMI Janelia Research Campus). pAAV-hSyn-CheRiff-eGFP (*61*) (Addgene plasmid # 51697) was a gift from Adam Cohen (Harvard). pTorPE-R-GECO1 (*62*) (Addgene Plasmid # 32465) was a gift from Robert Campbell (University of Alberta). pAdDeltaF6 (Addgene plasmid # 112867) and pAAV2/9n (Addgene plasmid # 112865) were gifts from James M. Wilson (University of Pennsylvania). All animal procedures were conducted following the protocols approved by the Animal Care and Use Committee at the University of Virginia and the University of Pittsburgh. Mice were hosted in temperature-controlled vivarium (∼ 23 °C) with a 12 h/12 h dark-light cycle and ∼50% humidity.

### Library construction and screening

The gene for the Rad50 zinc hook (AKGKCPVCGAELTD) was amplified from our previously reported ZIBG2 (*32*), and assembled via overlap extension PCRs with the fragments of the two cpmMaroon variants (cpmMaroon185-186 and cpmMaroon196-186) (*45*). The overlapped fragments were cloned into a modified pTorPE plasmid (*62*), which harbors a Twin-Strep-tag. Oligos with degenerate codons (NNK, where N = A, T, G or C, and K = G or T) were used to randomize the links between the zinc hooks and cpmMaroon. In addition, error-prone PCRs were used for random mutagenesis of the whole gene fragment. Library screening was performed similarly to a previously described procedure (*62*). Briefly, digital fluorescence images of bacterial colonies were taken before and after spraying 2 mM EDTA (a Zn^2+^ chelator) via a fine mist sprayer. Colonies showing the extreme fluorescent ratio changes were selected and used to inoculate 1 mL 2×YT medium supplemented with 100 μg/mL ampicillin and 0.2% (w/v) L-arabinose in 96-well deep-well plates at 250 rpm and 16 °C for 48 h. Cell lysates were then prepared from bacterial pellets using Bacterial Protein Extraction Reagents (B-PER). Fluorescence responses of the crude proteins to 100 μM Zn^2+^ or 100 μM EDTA were determined on a BioTek Synergy Mx microplate reader. Clones with large responses were chosen. Plasmids were prepared, sequenced, and used as templates in the further round of directed evolution.

### Protein purification and characterization

FRISZ in pTorPE was cloned into a modified pBAD plasmid harboring a Twin-Strep-tag. The resultant plasmid was used to transform E. cloni 10G competent cells (Lucigen). Protein expression was performed as described (*45*). Cell pellets harvested by centrifugation were next lysed by sonication. The Twin-Strep-tagged protein was purified via affinity chromatography using a StrepTrap HP column (GE Healthcare). Next, the eluant was subjected to size-exclusion chromatography through a HiLoad 16/600 Superdex 200pg column (GE Healthcare). Purified proteins were buffer-exchanged and concentrated in a HEPES assay buffer (150 mM HEPES, 100 mM NaCl, 0.5 mM TCEP, and 10% glycerol, pH 7.4) using Amicon Ultra Centrifugal Filter Units (10,000 Da molecular weight cutoff). Freshly prepared proteins were next diluted with the HEPES assay buffer mentioned above for *in vitro* assays on a monochromator-based BioTek Synergy Mx plate reader. Solutions containing 200 nM protein and 100 μM Zn^2+^ or 100 μM EDTA in the HEPES assay buffer were prepared. The emission wavelength was set at 670 nm to record the excitation spectra, and the excitation light was scanned from 450 to 630 nm. The excitation wavelength was fixed at 590 nm to record the emission spectrum, and the emission was scanned from 610 to 800 nm. For other endpoint measurements, the excitation and emission were set at 610 nm and 650 nm, respectively. Zn^2+^ titrations were performed by mixing the protein (a final concentration of 100 nM) with a series of HEPES assay buffers (*63*), which provided free Zn^2+^ concentrations ranging from 100 nM to 1 mM at 20 °C. The fluorescent intensity of each solution was determined and plotted as a function of the Zn^2+^ concentration. The data were then fitted to the Hill equation. To determine the metal selectivity of the protein, various metals (a final concentration of 100 μM) were added to the HEPES assay buffer containing 500 nM protein. The fluorescent intensity was recorded 10 min later.

### Determination of quantum yield, fluorescence lifetime, and extinction coefficients

For fluorescence measurements, the samples were diluted to have optical densities less than 0.09. Fluorescence quantum yields were determined using the absolute method with an integrating sphere instrument, Quantaurus-QY (Hamamatsu). In this measurement, the quantum yield (φ) was obtained at a set of excitation wavelengths changing from 500 and 590 nm with the step of 10 nm. For the Zn^2+^-free (100 μM EDTA) and Zn^2+^-saturated (100 μM Zn^2+^) samples, quantum yield did not systematically change in the region from 520 (or 510) to 590 nm, so the average of the corresponding measurements was accepted as a final value. Fluorescence lifetimes were measured on dilute solutions with a Digital Frequency Domain system ChronosDFD (ISS) appended to a PC1 (ISS) spectrofluorometer. Fluorescence was excited with a 635-nm laser diode (ISS, model #73296) through a 640/10 filter. The excitation was modulated with multiple harmonics in the range of 10−300 MHz. LDS 798 dye (Exciton) in ethanol with *τ*= 0.15 ns (*64*) was used as a lifetime standard to obtain the instrumental response function in each measurement. Fluorescence of the samples and standard were collected at 90° through HQ 705/100 and 695 LP filters to cut off all excitation light. The modulation ratio and phase delay curves were fitted to model functions corresponding to a single- or double-exponential fluorescence decay with Vinci 3 software (ISS). Only double exponential decay functions provided acceptable χ^2^ values, in the range of 0.59–0.73. The average fluorescence lifetime (calculated using intensity factors) is presented and used to calculate the extinction coefficient with the Strickler-Berg formula. Maximum extinction coefficients of the anionic forms (absorption at ∼ 606 nm) of the chromophore (ε_A_) were obtained using the Strickler-Berg formula that relates the extinction coefficient with fluorescence lifetime and quantum yield (*52*). To evaluate the extinction coefficient of the neutral form in both Zn^2+^-free and Zn^2+^-saturated states (absorption at 454 nm), we carried out a gradual alkaline titration of the samples and plotted the dependence of the neutral on anionic peak optical densities (*65*). The slope of this dependence is equal to the ratio of corresponding extinction coefficients. Neutral form extinction coefficient (ε_N_) was then found from this ratio and a known extinction of anionic form. The fractional concentrations of the anionic (ρ_A_) and neutral (ρ_N_) forms can then be calculated using initial optical densities (at pH 7.4) of the two corresponding absorption peaks and their extinction coefficients from Beer’s law. Here, indexes A and N correspond to the anionic and neutral forms of the chromophore, respectively

### Two-photon spectral characterization

Two-photon excitation spectra were measured as described (*66*). In the spectral shape measurement, Coumarin 540A in DMSO and LDS 798 in CHCl_3_:CDCl_3_ (1:2) were used as standards. A combination of 561 LP, 705/100, 770 SP, and 694 SP filters was used to block both the laser scattering and spurious fluorescence peaking at 550 nm and corresponding to a species absorbing near 530 nm. The cross-section σ_2_,A was measured at two close wavelengths, 1060 and 1064 nm. The measurement was performed using Rhodamine 6G (Rh6G) in methanol at λ = 1060 nm (for which σ_2_,_A_ = 10 ± 1 GM) (*66*) as a reference standard, and, independently, using Rhodamine B (RhB) in alkaline ethanol at λ = 1064 nm (for which σ_2_,_A_ = 13.3 ± 2.8 GM) (*51*) as an additional reference standard. The two-photon fluorescence signals of the sample and reference solutions in the same excitation and collection conditions were measured. We used a combination of the 770 SP, 705/100, and 561LP filters in the emission channel. The two-photon excitation spectra were then scaled to the obtained σ_2_,_A_ values to get the σ_2_,_A_ (λ) spectra. Both reference standards gave very similar results either for Zn^2+^-free or Zn^2+^-saturated samples. To obtain the two-photon excitation spectrum in units of molecular brightness, we independently measured ρ_A_, φ_A_ (the fluorescence quantum yield), and σ_2_,_A_ (λ) of the anionic form for FRISZ in Zn^2+^-free and Zn^2+^-saturated states and normalized the unscaled 2PE spectrum to the product F_2_ = ρ_A_•φ_A_ •σ_2_,_A_ (λ). The molecular brightness of the anionic form presented in the Table corresponds to the spectral maxima, λ_m_, for both states of the sensor.

### Construction of mammalian expression and viral packaging plasmids

The FRISZ gene was amplified from the pTorPE plasmid and inserted into a predigested pCMV(MinDis)-iGluSnFR plasmid between the Bgl II and Sal I restriction sites, resulting in a plasmid termed pDisplay-FRISZ. To construct the adeno-associated viral (AAV) transfer plasmid, a gene fragment encoding FRISZ, the N-terminal Igκ leading sequence, and the C-terminal PDGFRβ TM domain was amplified from pDisplay-FRISZ and inserted into a predigested pAAV-hSyn-CheRiff-eGFP plasmid between the BamH I and EcoR I restriction sites, resulting in a plasmid termed pAAV-hSyn-FRISZ-TM.

### Culture, transfection, and imaging of HEK 293T cells

HEK 293T cells were cultured and transfected as described (*32*). Fluorescence imaging was performed on a Leica DMi8 microscope equipped with a Leica SPE-II spectral confocal module and a Photometrics Prime 95B Scientific CMOS camera. Cells were washed with the Krebs-Hepes-bicarbonate (KRB) buffer (140 mM NaCl, 3.6 mM KCl, 0.5 mM NaH_2_PO_4_, 0.5 mM MgSO_4_, 1.5 mM CaCl_2_, 2 mM NaHCO_3_,3 mM glucose, 10 mM HEPES, pH 7.4) three times and a 40× oil immersion objective lens was used for imaging. To determine cellular localization, the confocal module was used. The far-red fluorescence was acquired with a 635 nm laser, and the emission was collected at 655-800 nm. In addition to pDisplay-FRISZ, a cytosolic EGFP (pcDNA3-EGFP) was expressed simultaneously for comparison. The green fluorescence was acquired with a 488 nm laser, and the emission was collected at 500-560 nm. To image the response of the indicator at the surface of HEK 293T cells, time-lapse experiments were performed under the wide-field condition using the Photometrics Prime 95B Scientific CMOS camera and a Cy5 filter cube containing a 628/40 nm bandpass excitation filter and a 692/40 nm bandpass emission filter. Cells were imaged every 10 s for a 15-min duration. Confocal images were also acquired between each addition of the different stimulating chemicals. To determine the binding affinity of FRISZ at the surface of HEK 293T cells, Zn^2+^ was added sequentially from low to high concentrations to gain 1-40 μM final concentrations, and cells were imaged every 1 min after thoroughly mixing the solution. To determine the half-time of the sensor for Zn^2+^ association and spontaneous dissociation, 200 μM Zn^2+^ was applied locally to the cells of interest, and time-lapse imaging with 15 ms exposure was performed. A glass micropipette with a ∼ 4 μm diameter tip was fabricated using a P-87 Flaming/Brown Micropipette Puller (Sutter), and a Picospritzer II microinjection dispenser was then used to deliver a buffered Zn^2+^ solution via the glass micropipette using a 3-ms puff.

### Preparation of Adeno-Associated Virus (AAV)

The AAV production was performed following a protocol from Rego *et al*. (*67*). The following plasmids (prepared from maxiprep) were used for AAV production: pAdDeltaF6, pAAV2/9n, and pAAV-hSyn-FRISZ-TM or pAAV-hSyn-mMaroon1-TM. Briefly, HEK 293T cells co-transfected with the plasmids were incubated in high-glucose Dulbecco’s Modified Eagle Medium (DMEM) supplemented with 4% fetal bovine serum (FBS), 0.1 M sucrose, and 10 mM HEPES for 96 h. Cells and media were collected and separated via centrifugation. The supernatant medium was filtered through a 0.45 μm polyethersulfone membrane filter, and viral particles in the supernatant were further precipitated using a solution containing polyethylene glycol 8000 and NaCl. The viral particles were resuspended in a cell lysis buffer (50 mM Tris, 150 mM NaCl, 2 mM MgCl_2_, pH 8.5). The pelleted cells were resuspended in the cell lysis buffer mentioned above and lysed by sonication. The sonication mixture was clarified by centrifugation and combined with the crude viral particles harvested from the medium. After being treated with 50 U/ml Benzonase nuclease, the sample was cleared via centrifugation and then further purified via gradient ultracentrifugation. Finally, the virus was collected and exchanged into Dulbecco’s phosphate-buffered saline (DPBS) containing 0.001% pluronic and 200 mM NaCl. The viral solution was aliquoted, flash-frozen, and stored at -80 °C. Viral titers were determined using a qPCR method provided by Addgene. The typical viral titers were ∼ 10^14^ genome copies per mL.

### Stereotaxic injections for *in vivo* imaging

C57/B6J mice between postnatal day (P) 28 and P36 were injected with the AAVs into the right auditory cortex as described previously (*47, 48*). Briefly, mice were anesthetized with isoflurane, and a craniotomy (∼0.4 mm diameter) was made over the temporal cortex (∼4 mm lateral to lambda). With a micromanipulator (Kopf), a glass micropipette containing the AAVs was inserted into the cortex 0.5–0.7 mm past the surface of the dura, and ∼500 nL of the virus was injected over the course of 5 min. Next, the scalp of the mouse was closed with cyanoacrylate adhesive. Mice were given carprofen 5 mg/kg (Henry Schein Animal Health) to reduce the pain associated with the surgery and monitored for signs of postoperative stress and pain.

### *In vivo* imaging preparation

About 21–28 days after the viral injections, mice were prepared for *in vivo* imaging as described previously (*47, 48*). Briefly, mice were anesthetized, head-fixed, and a craniotomy (∼1 mm diameter) over the A1 (∼4 mm lateral to lambda) was performed. To infuse the ZX1 into the auditory cortex, a micromanipulator (Siskiyou, Grants Pass, OR) was used to insert a glass micropipette backfilled with mineral oil and connected to a 5 μL glass syringe into the cortex at the edge of this craniotomy. The pipette contained ACSF and 100 μM ZX1. Next, mice were placed under the microscope objective in a sound- and light-attenuation chamber containing a calibrated speaker (ES1, Tucker-Davis Davis Technologies, Alachua, FL).

### *In vivo* wide-field imaging of the A1

After the *in vivo* imaging preparation, awake mice were used to image the sound-evoked activity in the auditory cortex in awake mice as described previously (*47, 48*). The Matlab-based Ephus program (*68*) was used to generate sound waveforms, and synchronize the sound delivery and image acquisition hardware. We presented sound stimuli (6-64 kHz broadband noise, 100 ms duration, 5 ms ramps at 30–80 dB SPL in 5 dB SPL increment level) while illuminating the skull with an orange LED (nominal wavelength of 590 nm, M590L4, Thorlabs). We imaged the change in FRISZ-TM or mMaroon1-TM emission with epifluorescence optics (Olympus U-MF2 filter cube: excitation filter-FF01-593/40 nm, emission filter-FF01-660/52 nm, and dichroic beam splitter-FF635-Di01, AVR optics, Rochester, NY) and a 4× objective (Olympus) using a cooled CCD camera (Rolera, Q-Imaging, Surrey, BC, Canada). Images were acquired at a resolution of 174 × 130 pixels (4× spatial binning, each pixel covered an area of 171.1 μm^2^) at a frame rate of 20 Hz in each mouse. A region of interest (ROI, 150–200 mm x 150–200 mm) over A1 was used to quantify sound-evoked responses. An average of the fluorescent intensity from all pixels in the ROI was derived. The sound-evoked change in fluorescence after the sound presentation (ΔF) was normalized to the baseline fluorescence (F_0_), where F_0_ is the average fluorescence of 1 s preceding the sound onset (for each pixel in the movie). ΔF/F responses from 6 to 8 presentations of the same sound level were further averaged. Response amplitude was the peak of the responses that occurred within one second of the sound onset. After the initial recording, the ZX1 solution (100 μM) was infused into the auditory cortex at a rate of 30 nL/min over 20 min. Next, the pump speed was reduced to 9 nL/min, and the sound-evoked responses were remeasured.

### Statistical and Data analysis

Microsoft Excel, GraphPad Prism, and Affinity Designer were used to analyze data and prepare figures for publication. Sample size and the number of replications for experiments are presented in figure legends. No statistical methods were used to pre-determine the sample size. Data are shown as mean and standard deviation (s.d.) or standard error (s.e.m), and the information is included in figure legends. Fiji (ImageJ) was used to analyze microscopic images. Imaging background was typically subtracted by setting the rolling ball radius to 50 pixels. ROIs were manually drawn, and average intensities were extracted for further analysis. When photobleaching was obvious, imaging stacks were corrected using an exponential decay fitting of baselines. When images were pseudocolored, the standard “Fire” lookup table was applied.

## Acknowledgments

We thank Drs. Xinyu Li, Yiyu Zhang, and Zefan Li for technical assistance and replicating some results. Research reported in this publication was supported by funding to H.A. (University of Virginia Start-up Package and NIH grants R01 DK122253, RF1AG077773, and R01 GM129291), T.T.(NIH grant R01 DC007905, NSF-IOS-1655480), and M.D. (NIH BRAIN grant U24 NS109107).

## Author contributions

HA conceived the project. TW performed indicator engineering, characterization *in vitro* and cultured cells, viral preparation, and initial tests in mice. SZ performed kinetics measurements and initial indicator tests in mice. XT refined the viral preparation protocol and prepared a batch of AAV. MD determined most photophysical parameters and performed 2-photon characterization. MK performed *in vivo* imaging. MK and TT designed the *in vivo* imaging experiments. TW, MD, and MK analyzed the data. HA, TW, MD, MK, and TT wrote the manuscript.

## Competing interests

Authors declare that they have no competing interests.

## Data and materials availability

The plasmids for pTorPE-FRISZ (Pladmid #176888), pDisplay-FRISZ (Plasmid #176887), pAAV-hSyn-FRISZ-TM (Plasmid #176886) and their sequence information have been deposited to Addgene. Other materials, research protocols, and supporting data are available in the main text or the supplementary materials, or from the corresponding author upon request.

## Notes

### Competing Interest Statement

The authors have declared no competing interest.

