## Supplementary for "A Genetically Encoded Far-Red Fluorescent Indicator for Imaging Synaptically-Released Zn^2+^"

**This PDF file includes:**

Figs. S1 to S4

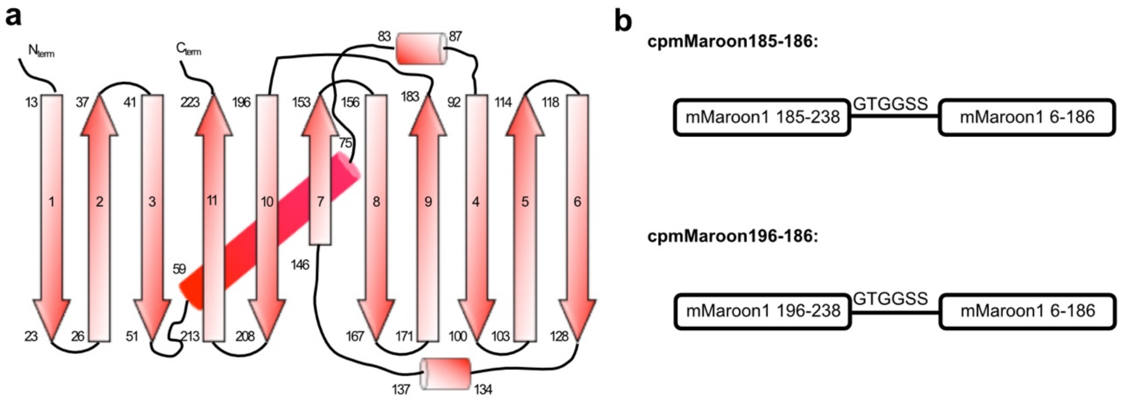


**Fig. S1. Illustration of the secondary structure of mMaroon1 and the primary structures of circularly permuted mMaroon1 variants.**

**(a)** Schematic representation of the secondary structure elements of mMaroon1. Cylinders represent α-helices, and arrows represent β-strands. The secondary structure elements are based on a mMaroon1 structure predicted by SWISS-MODEL using mCardinal (PDB: 4OQW) as the template. **(b)** Primary structures of cpmMaroon185-186 and cpmMaroon196-185 used in the indicator development. The residues 185-238 or residues 196-238 of mMaroon1 were linked to the residues 6-186 of mMaroon1 via a six-residue "GTGGSS" floppy linker.

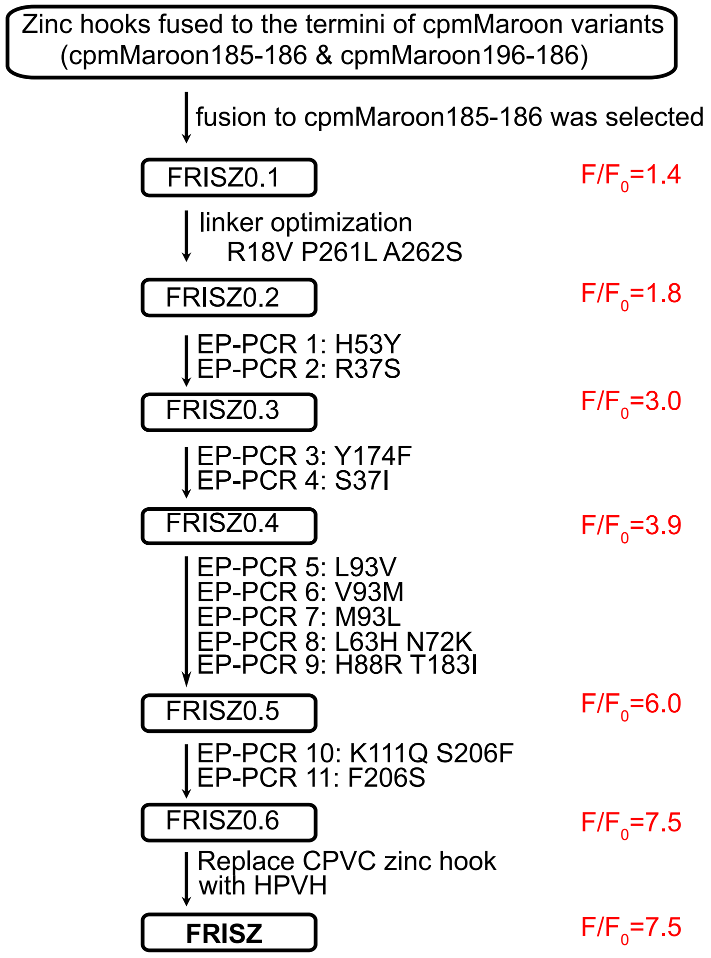


**Fig. S2. The process to engineer FRISZ.**

A flowchart is presented to illustrate our multi-step process to derive FRISZ. $F/F_0$ for each sensor generation is also shown.

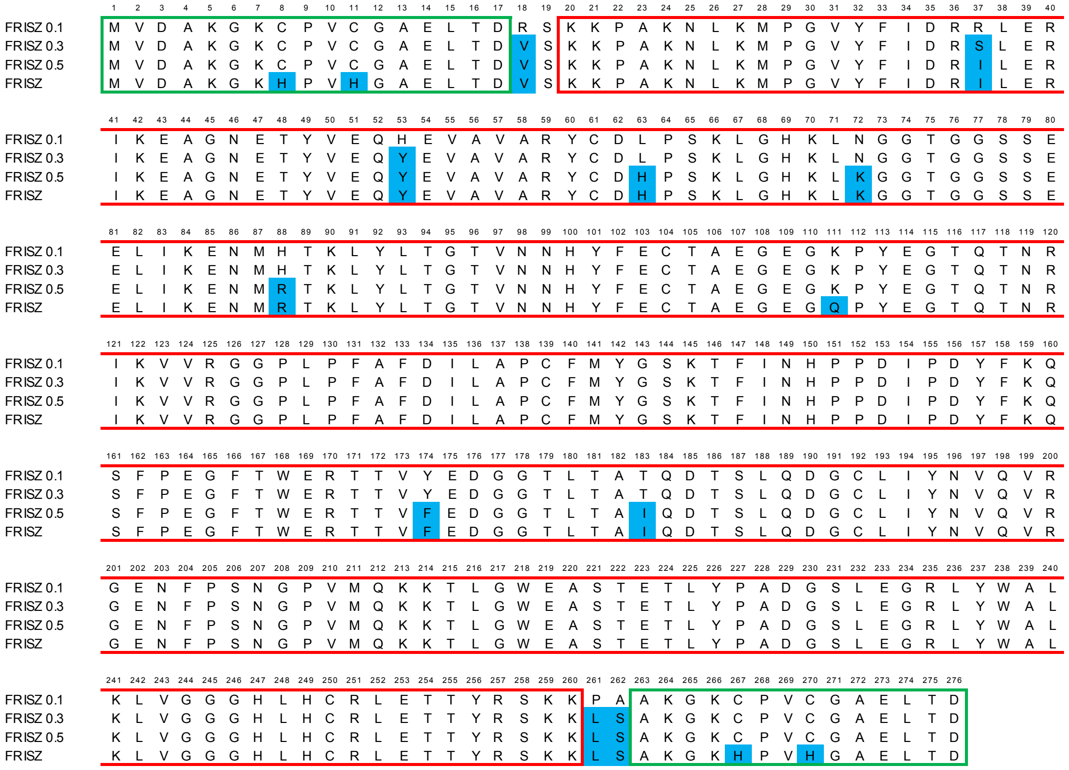


**Fig. S3. Sequence alignment of FRISZ with several earlier variants.**

The sequences derived from mMaroon1 and Rad50 zinc hooks are in red and green boxes, respectively. The mutations gained during the indicator engineering are shaded in blue.

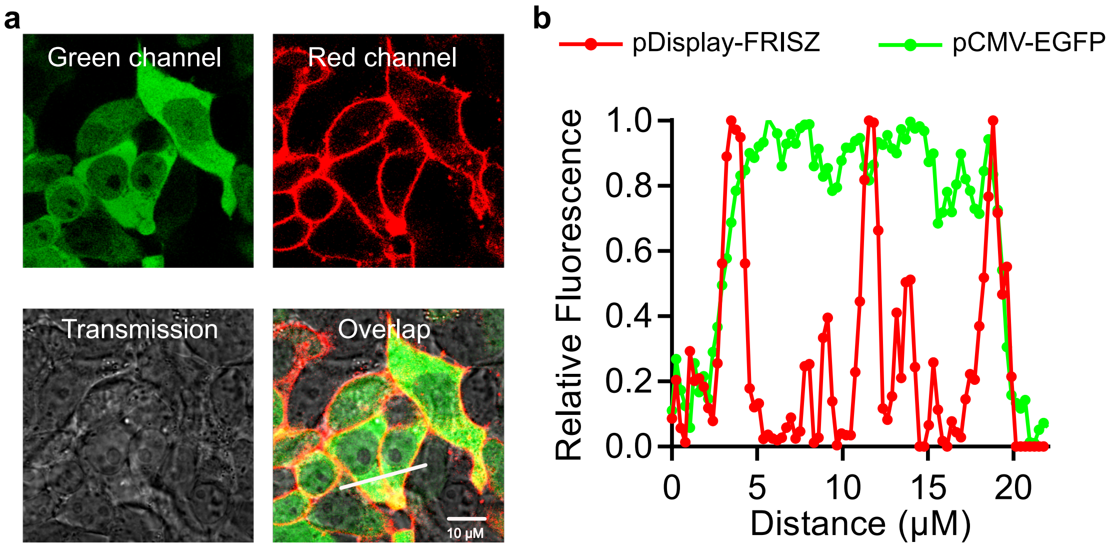


**Fig. S4. Characterization of pDisplay-FRISZ for membrane localization in mammalian cells.**

**(a)** Representative fluorescence images of HEK 293T cells co-transfected with pDisplay-FRISZ and pCMV-EGFP. For comparison purposes, EGFP was co-expressed as a whole-cell label. Scale bar, 10 µm. **(b)** Fluorescence intensity measured over the white line shown in the "overlay" image of panel a, further confirming the membrane localization of FRISZ in mammalian cells. These experiments were repeated three times with similar results using independent cultures.
